# Development of a cost-effective Alveolus-on-Chip for studying *Mycobacterium tuberculosis* infection

**DOI:** 10.64898/2026.04.29.721656

**Authors:** Nathalie Deboosere, Yoel Dagan, Aurélie Burette, Elise Delannoy, Sophie Salome Desnoulez, Elisabeth Werkmeister, Roxane Simeone, Priscille Brodin, Alexandre Grassart

## Abstract

We developed a cost-effective human alveolus-on-chip based on 3D printing molds (3DP-μLung) to study early events of *Mycobacterium tuberculosis* (*Mtb*) infection in a physiologically human relevant microenvironment. This organ-on-chip platform is compatible with advanced imaging and recreates the alveolar–capillary interface by co-culturing primary human alveolar epithelial cells, endothelial cells and macrophages. We show that epithelial-only models display limited susceptibility to *Mtb* infection, whereas the integration of macrophages significantly enhances infection levels of the alveolar barrier and supports intracellular bacterial replication. Quantitative imaging reveals that macrophages act as a permissive niche, promoting *Mtb* infection of both epithelial and endothelial compartments. This accessible organ-on-chip platform enables robust modeling of early events of host–respiratory pathogen interactions and provides a valuable tool for studying tuberculosis pathogenesis in human-relevant conditions. More broadly, it lowers technical and economic barriers to accelerate the adoption of organ-on-chip technologies for studying human specific infection.

**Summary:** A cost-effective human alveolus-on-chip enables physiologically relevant modeling of early host pathogen interaction and revealing a key role of macrophages in *Mycobacterium tuberculosis infection*

## Introduction

*Mycobacterium tuberculosis* (*Mtb*), the causative agent of tuberculosis (TB), remains one of the leading infectious killers worldwide, with an estimated 10.6 million new cases and 1.3 million deaths in 2022 (WHO, 2025). Despite being traditionally associated with immunocompromised individuals, including those with HIV infection, TB also imposes a substantial burden on immunocompetent populations, particularly in low- and middle-income countries where transmission is endemic. Pulmonary TB is the most common and contagious form of the disease, initiated when inhaled bacilli reach the alveolar space and encounter resident immune and epithelial cells (Torrelles & Schlesinger, 2017; Leon-Icaza et al, 2025). Following phagocytosis by alveolar macrophages, *Mtb* can subvert host antimicrobial defenses, establish an intracellular niche, and eventually disseminate to other immune and parenchymal cells (Warner et al, 2025).

Traditional *in vitro* models for *Mtb* infection have largely relied on two-dimensional cultures of macrophages or epithelial cells, as well as immortalized monocytic cell lines such as THP-1 and U937 (Kilinç et al, 2022). Although these systems are suitable for single cell mechanistic studies, including high-throughput screening campaigns, they fail to recapitulate key structural and physiological features of the human alveolus, including three-dimensional alveolar–capillary interface, coordinated innate immune responses and other tissue scale characteristic such as air–liquid interphase. Animal models, particularly murine systems, have been indispensable in elucidating aspects of TB immunopathology (Flynn et al, 2015; Warner et al, 2025). However, significant interspecies differences in lung physiology, immune responses, and granuloma formation limit their translational relevance to human disease.

Recent advances in organoid and organ-on-chip (OOC) technologies have opened new avenues to model TB infection in a human-relevant context. Organoid-based systems have provided valuable insight into *Mtb* pathogenesis and host responses (Alcaraz et al, 2022; Iakobachvili et al, 2022). Human lung on chip have further advanced the field by reconstituting complex alveolar–capillary barrier including mechanical (Huh et al, 2010) and immunological cues (Mishra et al, 2023; Luk et al, 2025). By culturing on opposite sides alveolar epithelial cells and lung microvascular endothelial cells within microfluidic channels separated by a porous membrane, many lung-on-chip models enabled the recreation of the alveolar–capillary interface under controlled flow, air–liquid interphase, and cyclic mechanical stretch (Jain et al, 2018; Li et al, 2019). Such new biomimetic capacities have been successfully applied to study the infection biology of *Mtb* (Thacker et al, 2020; Mishra et al, 2023), as well as other pathogens such as *Shigella, Escherichia coli*, *Aspergillus fumigatus*, *Staphylococcus aureus*, and influenza virus (Huh et al, 2010; Grassart et al, 2019; Huang et al, 2021; Bai et al, 2022; Mishra et al, 2023). Despite their promises, these new alternative models (NAMs) still face significant barriers to implementation in wet laboratories. These include the technical complexity of the microfluidic device fabrication using heavy lithography and microfabrication techniques, the biological intricacy of human cell co-culture and the variability in device design between platforms. Although commercial solutions address many of these limitations, their high cost remains an important obstacle to widespread adoption in academic labs.

Here, we report the development of a rapid and cost-effective human alveolus-on-chip model based on a facile fabrication approach using 3D printed molds as recently developed for gut on chip (Delannoy et al, 2025). This interoperable and versatile method enables robust production of OOC compatible with advanced imaging microscopy. We present and validate a methodology combining primary human alveolar epithelial cells, microvascular endothelial cells and macrophage cells co-cultured on the new open-sourced OOC platform, named 3DP-μLung, to reconstitute a physiologically relevant human alveolar barrier. We then demonstrated the capacities of this system by investigating the host–pathogen interactions of *Mtb* within this complex biomimetic microenvironment and dissected the early events of *Mtb* infection, during the first five days. Using confocal imaging, we characterized cellular tropism in response to *Mtb* infection, highlighting the benefits of OOC systems as a physiologically relevant model to advance mechanistic understanding of TB pathogenesis.

## Results

### Development of 3DP-µLung chip, a cost-effective Alveolus-on-Chip (AoC) optimized for imaging

We developed an in-house alveolus-on-chip (AoC), named 3DP-µLung platform and designed to be cost-effective, easily manufacturable in a biology laboratory. To promote robustness and cross standardization between labs, we adopted the widely established chip configuration, which comprises two microfluidic channels separated by a porous membrane (Huh et al, 2010). However, geometry and dimensions of our microfluidic device have been specifically optimized to enhance compatibility with high-speed confocal imaging (Fig 1A). In particular, the working distance between the lens and the porous has been reduced to avoid the need of a long-distance objective and enables the use of conventional air-dry objective for rapid confocal microscopy. Briefly, the bottom side of the chip is supported by a thin layer of PDMS i.e. 100 µm. In addition, the total length of the channels was halved as compared to published lung on chip based on similar design (Huh et al, 2010), thereby decreasing the amount of human primary cell required at seeding stage without compromising barrier formation.

**Figure 1.**
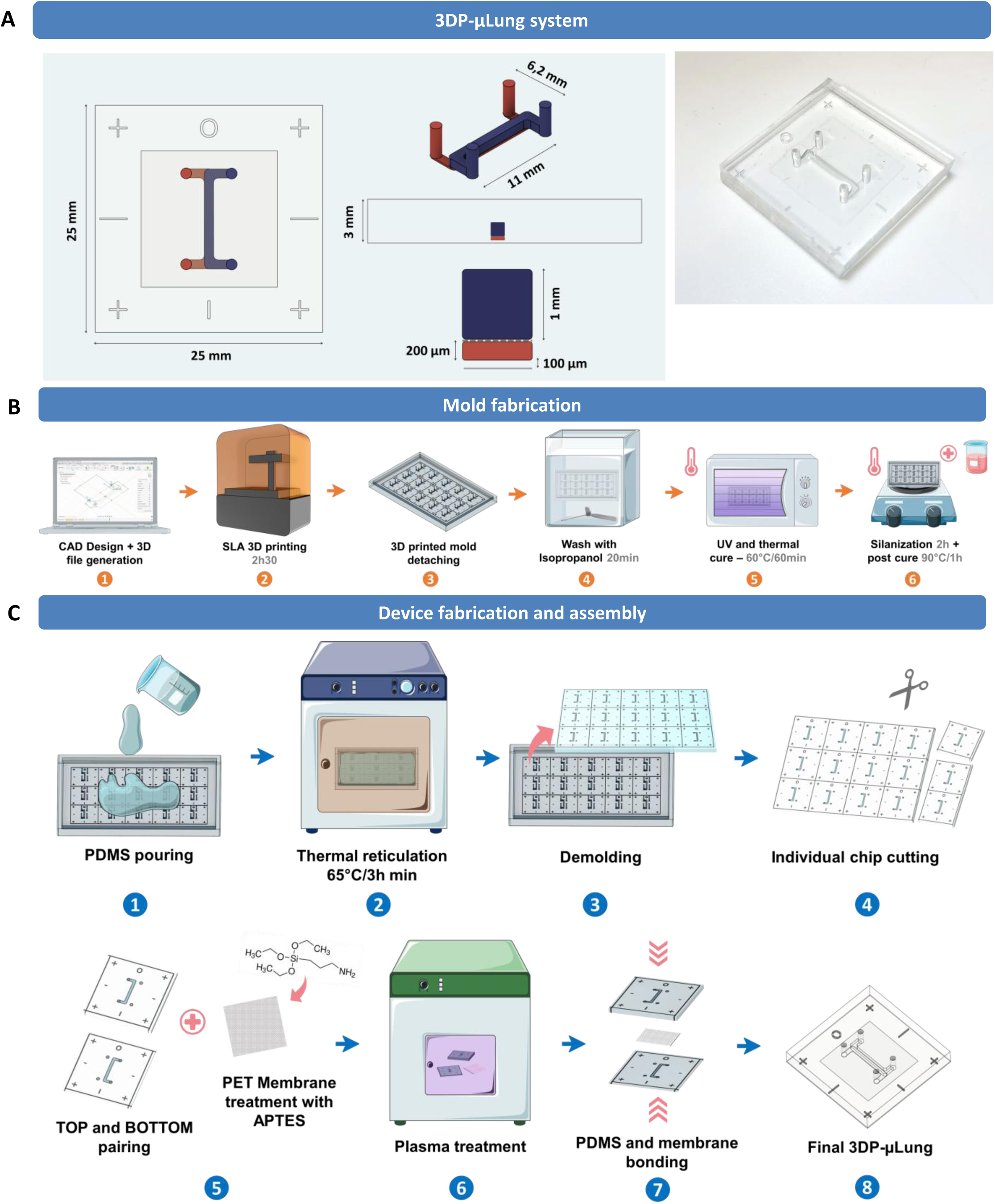
In-house 3DP-µLung microdevice. **(A)** Schematic view of the 3DP-µLung model. (Left) 3D view, top view, vertical cross-section. The top channel is in red and the bottom channel in blue. (Right) Photograph of the assembled 3DP-µLung chip **(B)** Fabrication process for the 3D printed molds and post treatment steps. **(C)** Assembly steps for the 3DP-µLung chip device.

Device fabrication was adopted from our previously established protocol for 3DP-μGUT model (Delannoy et al, 2025). Briefly, using a commercial high-resolutive 3D printer combined with soft lithography approach, we manufactured two molds for the lower and the upper microfluidic channels (Fig 1B), each mold containing up to fifteen identical chip units. To prevent PDMS uncuring and undesired adhesion of the silicon to the mold, post-curing and silanization steps were performed before PDMS casting (Fig 1B and C). Eventually, the 3DP-µLung device consists of the bonding of top and bottom parts with a purchased track-etched PET membrane in between (Fig 1C). Overall, this process takes about one to two day and reduces potential economic barrier linked to organ-on-chip technology.

### Alveolar-capillary interface formation and alveolar cell differentiation

Next, to reconstruct the alveolar-capillary barrier in the 3DP-μLung chip, an extracellular matrix (ECM) composed of fibronectin and collagen IV, both physiological components of human basement membranes, was coated on both sides of the porous membrane the day prior seeding of human primary alveolar epithelial cells (“HPAEpiC”) and lung microvascular cells (“HuLEC”) (Fig 2A). In a standardization effort, HPAEpiC and HuLEC cells were obtained from a conserved commercial source as previously published (Thacker et al, 2021; Bai et al, 2022; Mishra et al, 2023; Ektnitphong et al, 2024; Man et al, 2024). After four hours, most cells initially loaded in channels appeared fully spread across the surface of the porous membrane (Fig 2B). HPAEpiC are composed of alveolar type I (AT1) and type II (AT2) cells. Cell seeding density was optimized by transmitted light microscopy to achieve a homogeneous cell monolayer, with an estimated cell to cell spacing of approximately the radius of a single cell. All maturation steps of the alveolar-capillary interface were followed daily under phase contrast microscopy (Fig 2C). Cell monolayers reached confluence after 2 days of culture in static conditions. Then, cells exposed to a continuous flow at 60 μl/h in both channels for 4 days. Dexamethasone was added to the apical cell culture medium to enhance alveolar epithelial barrier function (Bai et al, 2022). At day 6, the apical medium was chased out and HPAEpiC were cultured in an air-liquid interface (ALI) to better mimic the alveolar environment, while a continuous flow through the lumen of the lower vascular channel was maintained. The culture was exposed in ALI for one day, after which the AoC was returned to a liquid-liquid interface (LLI) during macrophage co-culture and/or Mtb infection; the infection then proceeded for up to five days.. We validated the structuration and morphology of cells under the physiological flow. We observed a confluent and tight cell barrier by microscopy and visualized a homogenous distribution of cells across the entire chip within 13 days (Fig 2D). High-resolution confocal microscopy enabled the 3D reconstitution of the epithelium and endothelium barrier on either side of the membrane within the chip (Fig 2E). A cross-sectional image from the projection of all orthogonal sections of the obtained alveolar-capillary barrier confirmed the formation of a continuous barrier (Fig 2F). We also confirmed no significant cytotoxic cell death and no loss of cell viability on day 13 using live/dead assay (Fig 2G). Based on image analysis, we determined the number of cells grown in a matured AoC by detecting and counting nuclei intensity (DAPI). A classification between the two cell types was based on cells height in the slice thickness. The approximate total cell counts across the entire surface area of the chip membrane, inferred from confocal imaging, were 5.10^3^ for HuLEC and 2.10^4^ for HPAEpiC.. The characterization of the barrier was further achieved using known markers of barrier maturation by immunofluorescence and qRT-PCR analysis (Thacker et al, 2021; Ektnitphong et al, 2024). Staining for vascular endothelial-cadherin (VE-cadherin) revealed a flat HuLEC monolayer and cells colonized the entire channel on the endothelial side (Fig 3A). While the majority of the epithelium exhibit a monolayer structuration, we observed the existence of rare sporadic 3D protrusion-like architectures (Figs 3B and C). Furthermore, we confirmed by both immunostaining and qRT-PCR (Fig 3C and D) that the alveolar epithelium includes both type I (AT1) and type II (AT2) alveolar cells. For immunostaining, HT1 and HT2 were used as markers to detect AT1 and AT2 cells, respectively.. By day 8, a significant fraction of AT1 cells was present relative to AT2 cells, with a subset of cells exhibiting an AT2-to-AT1 transitional phenotype.(Fig 3C), as previously reported (Ektnitphong et al, 2024). Analysis of epithelial cell gene expression on uninfected chips revealed no extensive changes in mRNA expression for type I (AQP5, PDPN, CAV1) and type II (ABCA3, SFTPC) markers overtime (Fig 3D). Therefore, mature AoC retained its differentiation in AT1 and AT2 cells, which is consistent with previously reported results for other Lung-on-Chip models using these HPAEpiC (Thacker et al, 2021; Ektnitphong et al, 2024). Finally, the alveolar-capillary barrier permeability was evaluated by measuring transport of 70 kDa FITC–dextran across the alveolar barrier. As a control we used a non-fully matured chip, for which gaps in the monolayer were visualized upon inspection by phase contrast microscopy. The measured values of apparent permeability (Papp) showed a robust barrier formation as compared to the non-matured control condition (Fig 3E). Overall, our results confirmed that an effective alveolar barrier was established under these culture conditions.

**Figure 2.**
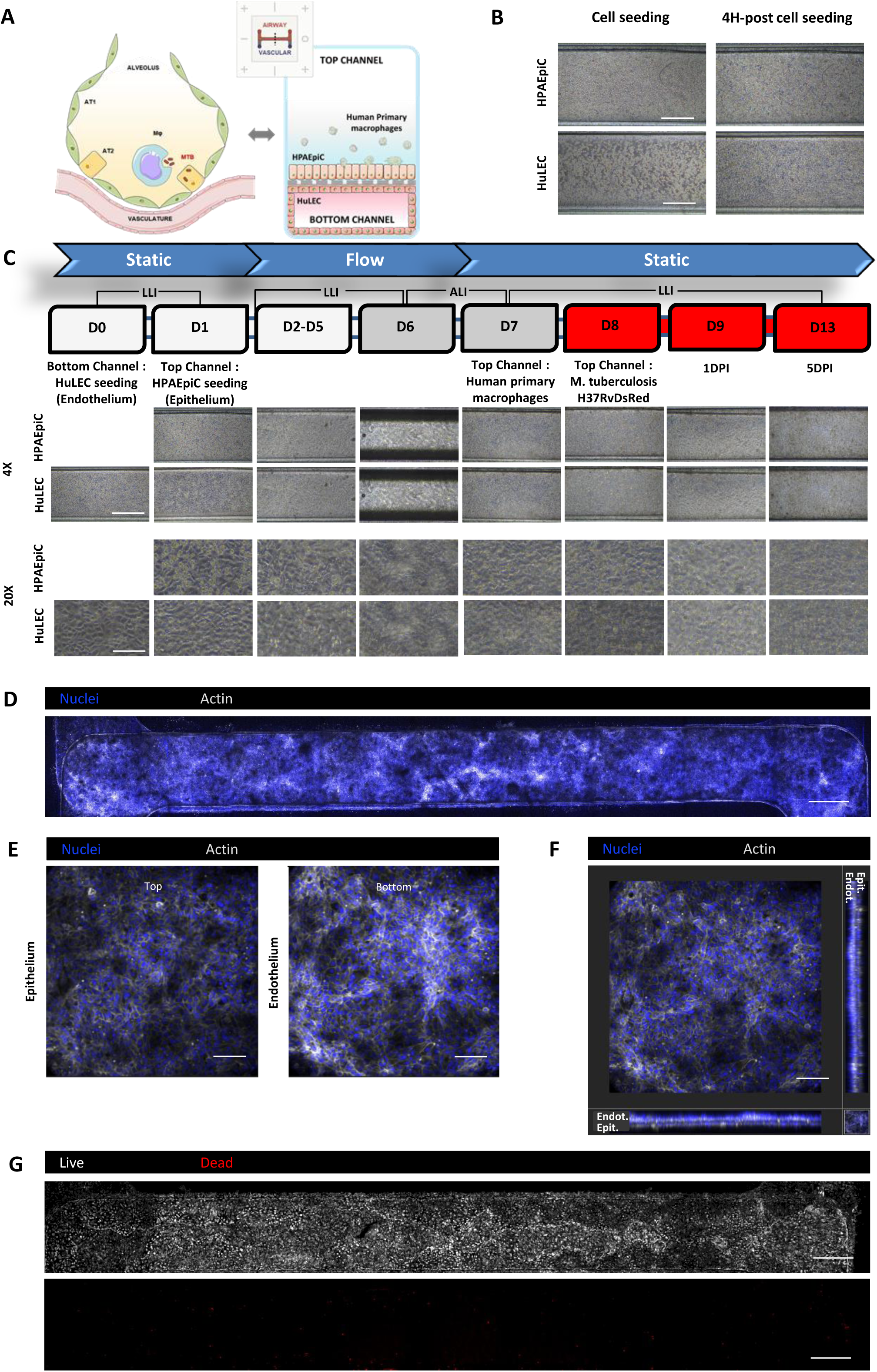
In-house 3D-Lung Alveolus-on-Chip (AoC) model, corresponding to an alveolar interface reconstituted with HPAEpiC and HuLEC cells. **(A)** Schematic vertical cross-section of an alveoli and AoC model. **(B)** Bright-field images of typical cell densities, obtained immediately after cell seeding and after 4-hours incubation. Scale bar = 500 μm. **(C)** Bright-field images of HuLEC and HPAEpiC co-culture inside the device during AoC maturation. Scale bar = 500 μm (4X) & 100 μm (20X). **(D, E, F, G)** Immunofluorescence staining of an uninfected matured AoC on day 13. Nuclei (Blue), Actin (White) except in G where live cells are White and dead cells are in Red **(D)** Maximum intensity projections of Z-stack slices of the entire top and bottom channels. Scale bar = 500 μm **(E)** 3D-images of z-slices, corresponding to HPAEpiC (Top) and HuLEC (Bottom) layers. Scale bar = 100 μm. **(F)** Maximum intensity projections of z-slices of the entire top and bottom channels for Live/Dead assay. Scale bar = 100 μm. **(G)** 3D view of the Z-stack images. Scale bar = 500 μm.

**Figure 3.**
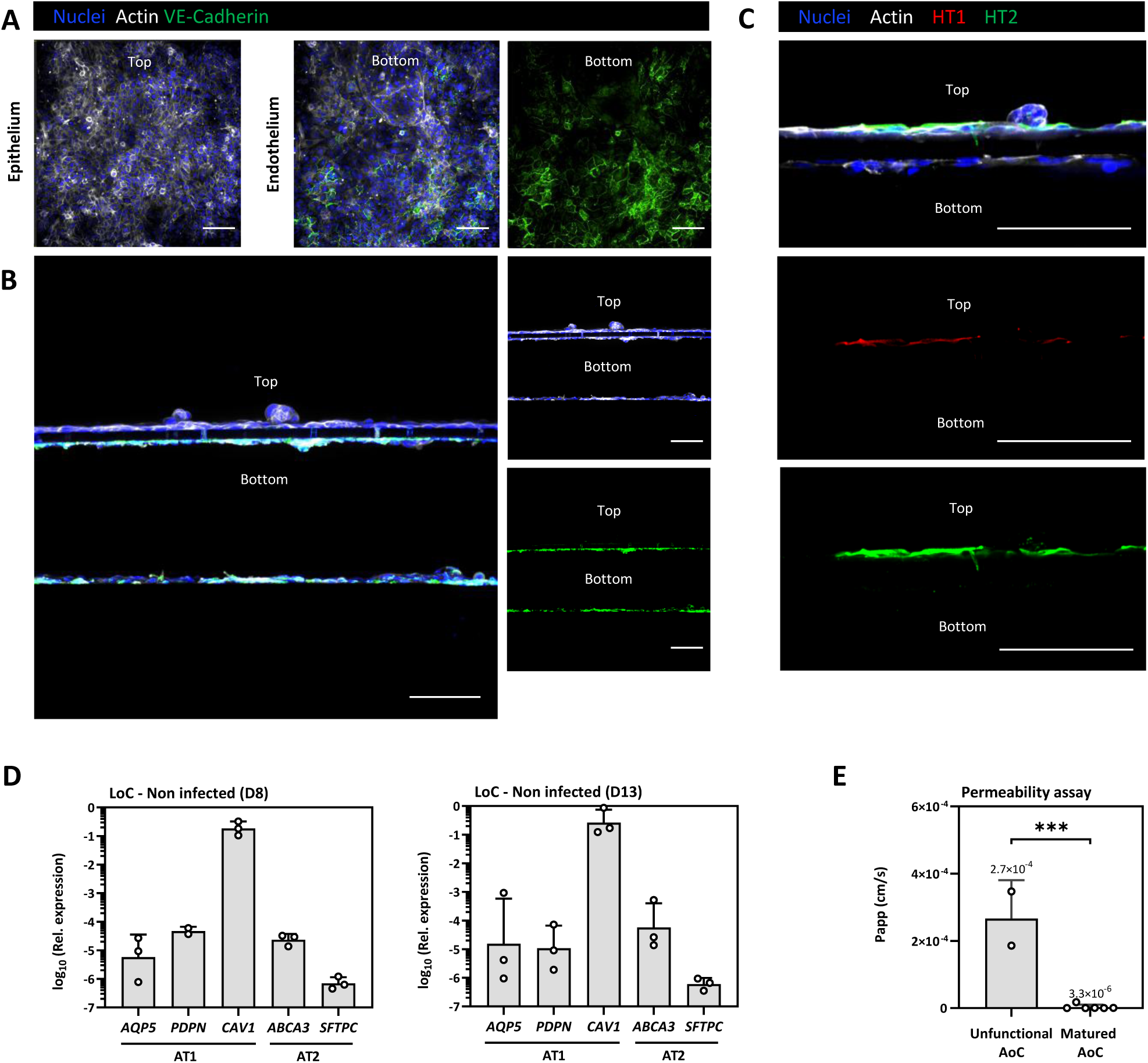
Characterization of alveolar barrier. **(A, B, C)** Immunofluorescence staining of an uninfected AoC on day 8. **(A)** 3D-images of z-slices, corresponding to HPAEpiC (Top) and HuLEC (Bottom) layers. Nuclei (Blue), Actin (White) and VE-Cadherin (Green), Scale bar = 100 μm. **(B)** Transversal sections, showing 3D-organization of HPAEpiC layer in top channel and HuLEC monolayers in the bottom channel and **(C)** confirming expression of alveolar epithelial type I (AT1) and type II (AT2) markers (HT1+ and HT2+, respectively) in epithelial cell barrier. Nuclei (Blue), Actin (White), HT1 (Red) and HT2 (Green), Scale bars = 100 μm. **(D)** qRT-PCR characterization for type I & II alveolar epithelial markers from uninfected AoCs extracted on day 8 (n=3) and day 13 (n=3). Expression of AT1 and AT2 markers was normalized to GAPDH levels. **(E)** Functional alveolar barrier validation by dextran permeability assay in the AoC in flow conditions. Uninfected AoC after 7-day post maturation (n=6) were compared to unfunctional AoC (n=2), showing no cell barrier integrity on day 7 and day 13. Unpaired t test was used to compared unpaired data. *P≤0.0005, ns = nonsignificant.

### An Alveolus-on-Chip model of *Mtb* lung infection

Following maturation, the AoC model was used to characterize early *Mtb* infection dynamics via microscopy and image analysis.To do so, we used a multiplicity of infection (MOI) of 5, which is comparable to conventional in vitro 2D experiments, to achieve sufficient infection of HPAEpiC cells within 4 hours (Fig. S1). Epithelial cells were exposed to DsRed-expressing *Mtb*, and the infection was monitored at 1 or 5 days post-infection (dpi). To quantify *Mtb* infection via microscopy, a maximum intensity projection was generated for each field of view, superimposing the epithelial and endothelial layers of the infected AoC. Quantitative image analysis was then conducted on the H37Rv-DsRed signal (Fig 4A). The cross-sectional image of the AoC showed that *Mtb* was mainly detected in the epithelial cell layer (Fig 4B), signifying successful epithelial infection events. Then, a dedicated image analysis script (Fig. S2) was developed using Imaris 3D software to quantify total cell counts (Fig. 4C), the bacterial load within the alveolar barrier (Fig. 4D), and the subsequent intracellular growth of *Mtb*. (Fig 4E). We first compared the total cell counts per ROI at 1 and 5 dpi, demonstrating that confluent monolayers of HPAEpiC and HuLEC were well-established after 9 and 13 days, respectively. Furthermore, no significant barrier damage was observed following *Mtb* infection under these conditions (Fig. 4C). (Fig 4C). We found that epithelial cells were twice as numerous as endothelial cells, which is likely due to differences in cell morphology; AT2 cells are more cuboidal and compact, whereas endothelial cells are flatter and more elongated. We observed a total infection rate of 0.5% at one day post-infection (Fig. 4D), comparable to the rate observed in conventional 2D culture plates (384-well) after 4 days of infection when cell confluency was reached (Fig. S1). This highlights the low infectivity of human alveolar epithelial cells, consistent with a previous report (Luk et al., 2025). By day 5 post-infection, we observed a non-significant increase in the infection rate to 1%. Similarly, no significant bacterial replication was observed within the alveolar barrier model after 5 days (Fig. 4E). Taken together, our results suggest that Mtb infectivity and growth remain low within a human alveolar-capillary interface, consistent with observations in other models (Thacker et al, 2020; Luk et al, 2025).

**Figure 4.**
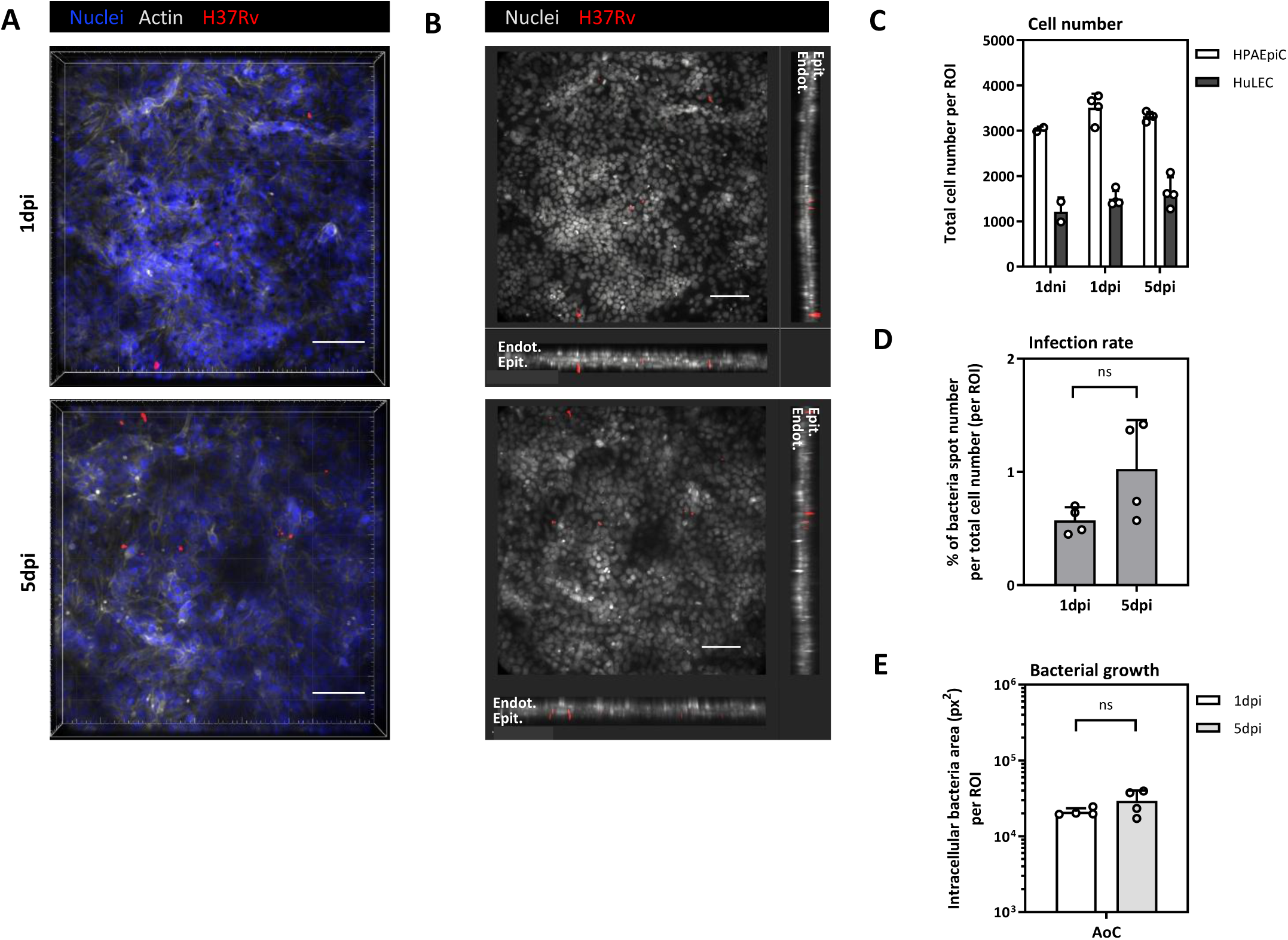
*Mtb* infection in 3D Alveolus-on-Chip. **(A, B)** Immunofluorescence imaging of infected AoCs at 1dpi and 5dpi, used for the determination of H37Rv-DsRed infection inside the alveolar barrier model. **(A)** 3D-images of z-slices, and **(B)** maximum intensity xy projections and extended orthogonal xz and yz cross-sections. Nuclei (Blue), Actin (White) and H37Rv (Red). Shown are representative images of four independent chips from two independent experiments, using a 3D image analysis software (IMARIS v10, Oxford Instruments). Scale bar= 100 μm. **(C, D, E)** Quantitative image analysis of non-infected and infected AoC on day 9 (1dpi and 1dpi, respectively) and infected AoC on day 13 (5dpi). **(C)** Cell numbers, **(D)** percentage of bacteria spot number per total cell number, and **(E)** bacterial growth, determined from Region of Interest (ROI) of 3D-culture model. Data are presented as mean ± SEM (n=4). Non-parametric Mann-Whitney test was used to compared unpaired data. *P≤0.05, ns = nonsignificant.

### Effects of macrophages on intracellular *Mtb* colonization within the alveolar barrier

As macrophages are a well-established niche of infection (Cohen et al, 2018; Ahmad et al, 2022; Bo et al, 2023; Sankar & Mishra, 2023), we investigated *Mtb* infectivity and growth within a alveolar barrier incorporating human macrophages. Briefly, these cells were derived from peripheral blood cells and engrafted into the AoC from the epithelial interface for twenty-four hours prior to infection. In agreement with alveolar physiology and previous studies, we chose a macrophage-to-HPAEpiC ratio of 1:10 (10%)(Weibel, 2015; Thacker et al, 2020). To facilitate direct monitoring of the macrophages within the alveolar barrier, cells were pre-stained with carboxyfluorescein succinimidyl ester (CFSE). This approach allowed us to visualize macrophage adherence via fluorescence microscopy (Fig. 5A) and to observe their dynamics using live-cell spinning disk confocal microscopy. (Video S1). We further validated this approach using Alexa Fluor 488-conjugated anti-CD68 labeling. Immunofluorescence imaging of the macrophage-containing AoC (M-AoC) at 1-day post-infection revealed that CFSE-labeled adherent macrophages and intracellular DsRed-expressing bacteria were distributed along the apical channel (Fig. 5A). These observations confirmed successful macrophage adhesion, integration, infection, and homogeneous distribution. Subsequently, we developed an image analysis pipeline to assess the impact of macrophages on *Mtb* replication within the alveolar barrier. During pulmonary infection, macrophages are generally considered the primary host cells; however, alveolar epithelial cells have also been identified as a critical initial contact site for the pathogen. (Ryndak & Laal, 2019; Moule & Cirillo, 2020; Sankar & Mishra, 2023; Olmo-Fontánez et al, 2024). Here, whole-mount imaging of the chip (Fig. 5A) and frontal cross-sections of the infected M-AoC at 1 dpi (Fig 5B) showed effective infection of the “immunocompetent” alveolar barrier. We observed partial colocalization of Mtb, and with the macrophage population (Fig 5A-B). We detected large clumps of *Mtb* bacteria scattered throughout the alveolar barrier after 5 days of infection within M-AoC (Fig 5C and D). Resolution of our imaging system did not allow us to determine whether these clumps represent precisely cord-like structures (Figs 5C and S3). Furthermore, we confirmed that cell diversity remained stable between 1 and 5 dpi (Fig. 5E). This indicates the sustained presence of both HPAEpiC and HuLEC layers at both time points, regardless of whether macrophages were present. Interestingly, we also observed that macrophages co-culture within the alveolar barrier led to *Mtb* infection of both cell types, epithelial cells and macrophages (Fig S3). Cross-sectional images of the AoC revealed that *Mtb* and macrophages were predominantly localized within the epithelial cell layer (Fig 5D). However, our quantitative image analysis (Fig S2) further revealed a significant increased number of infected macrophages at 5 dpi as compared to 1 dpi (Fig 5F). Moreover, we notably observed that sometimes macrophages migrated through the membrane to endothelial compartment at 5 dpi (Figs 5D and S3), a phenotype also reported in mouse lung on chip model (Thacker et al, 2020).

**Figure 5.**
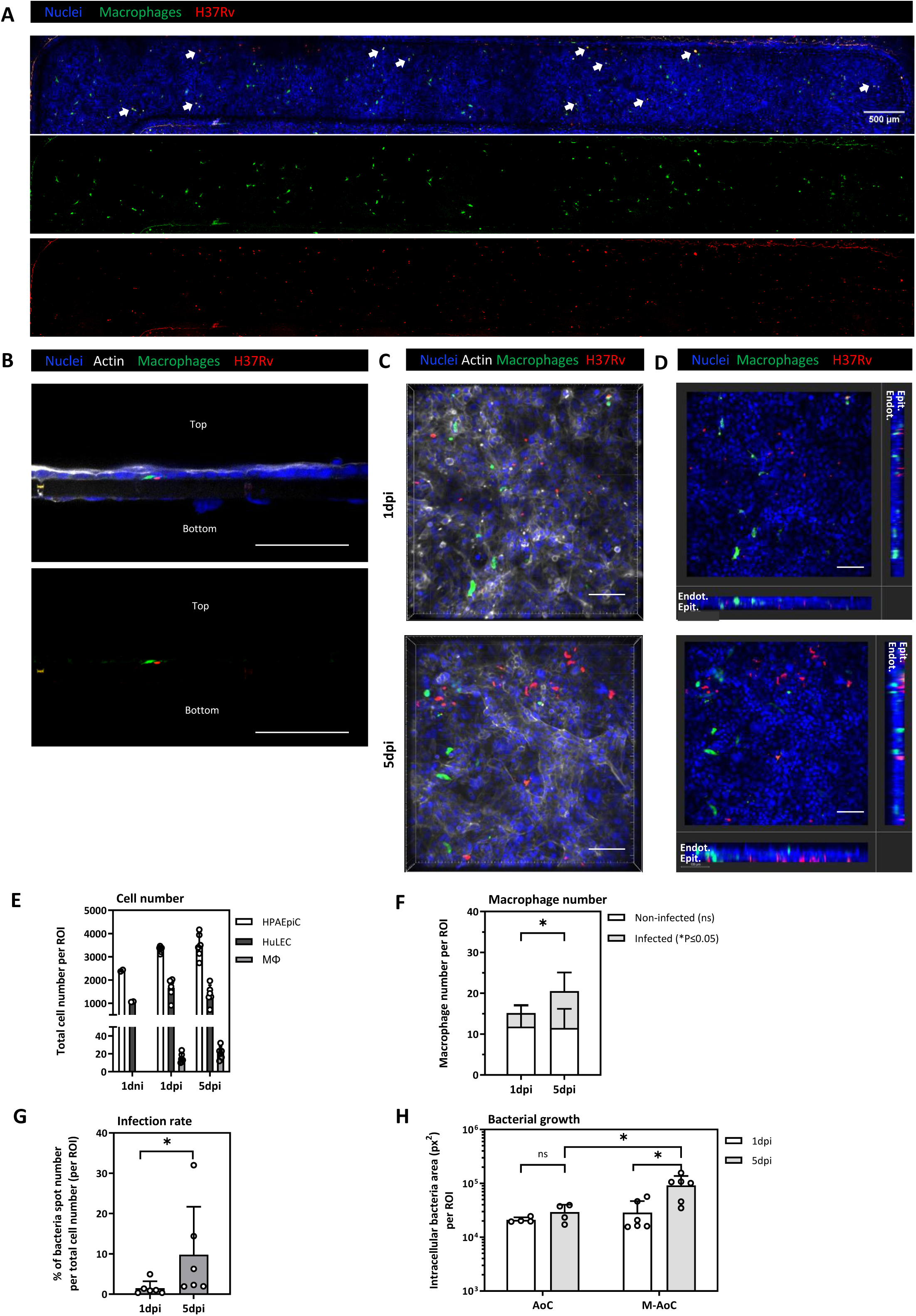
*Mtb* infection in Alveolus-on-Chip co-cultured with human macrophages (M-AoC). **(A, B)** Immunofluorescence staining of infected M-AoC on day 9 (1 dpi). (A) Top view, corresponding to complete imaging of chip, obtained by maximum intensity projections of z-slices of the entire top and bottom channels of infected M-AoC, highlighting co-localized bacteria and macrophages (white arrows). Nuclei (Blue), H37Rv (Red) and macrophages (Green), Scale bar= 500 μm. **(B)** Frontal cross-sections of infected M-AoC. Nuclei (Blue), H37Rv (Red), Actin (White) and CFSE/CD68 FITC (Green), Scale bar=100μm **(C, D)** Immunofluorescence imaging of infected M-AoC at 1dpi and 5dpi, used for the determination of H37Rv-DsRed infection inside the alveolar barrier model. Nuclei (Blue), H37Rv(Red), Actin (White) and CFSE/CD68 FITC (Green), **(C)** 3D-images of z-slices and **(D)** maximum intensity xy projections and extended orthogonal xz and yz cross-sections. Shown are representative images of six independent chips from two independent experiments, using a 3D image analysis software (IMARIS v10, Oxford Instruments). Nuclei (Blue), H37Rv (Red), Actin (White) and macrophages (Green). Scale bar = 100 μm. **(E, F, G, H)** Quantitative image analysis of non-infected and infected M-AoC on day 9 (1dni and 1dpi, respectively) and infected M-AoC on day 13 (5dpi). **(E)** Cell numbers, **(F)** numbers of non-infected and infected macrophages, **(G)** percentage of bacteria spot number per total cell number and **(H)** bacterial growth, determined from Region of Interest (ROI) of 3D-culture model. Data are presented as mean ± SEM (n=6). Non-parametric Mann-Whitney test was used to compared unpaired data. *P≤0.05, ns = nonsignificant.

Because bacteria segmentation at low magnification (10x to 20x air objectives) is challenging in organ-on-chip models, the number of detected bacteria may be underestimated due to bacterial clustering. However, our quantitative analysis revealed that approximately 10% of total cells of the alveolar barrier were infected at 5 dpi (Fig 5G). This level of infection is consistent with previously reported in vivo observations in mice and in mouse organ-on-chip models (Thacker et al, 2020). Finally, we interrogated if macrophage co-culture impacts intracellular *Mtb* growth. To do so, we analyzed the bacterial surface area by quantitative image analysis and observed a significant active replication after 5 days post infection in immunocompetent alveolar barrier as compared to alveolar barrier without macrophages (Fig 5H). Altogether, our results confirm that macrophages are a physiologically relevant *Mtb* infectious niche but also reveals that macrophages play a key and wider role in the early steps of *Mtb* infection by sustaining active infection other cell types of the alveolar barrier. Overall, these results demonstrate that our AoC model supports a physiologically robust human alveolar interface, effectively recapitulating key features of the early stages of pulmonary tuberculosis.

## Discussion

We successfully established a human alveolus-on-chip platform to study early events of *Mtb* infection using advanced microscopy-based approaches and quantitative analysis. In recent years, numerous OOC platforms have been developed to recapitulate structural and functional complexity of various human tissue or organ, including the lower and upper respiratory tract (Huang et al, 2021; Thacker et al, 2021; Bai et al, 2022; Francis et al, 2022; Ingber, 2022; Sengupta et al, 2022; Van Riet et al, 2022; Mishra et al, 2023; Ektnitphong et al, 2024; Man et al, 2024; Luk et al, 2025). Here, our lung on chip relies on the proposed 3DP-μLung platform, specifically designed to facilitate a broad implementation of OOC by reducing the fabrication cost and by simplifying the fabrication process. In contrast to conventional cleanroom-based lithography microfabrication (Huh et al, 2013), 3DP-μLung is fabricated using 3D printed molds. This proven approach unlocks rapid prototyping (hours versus days), reduces fabrication cost and facilitates transfer of the technology between laboratories (Novak et al, 2018; Shrestha et al, 2019; Byrne et al, 2023; Delannoy et al, 2025). Importantly, the chip geometry was optimized to monitor early host-pathogen interactions at the alveolar interface using high-resolution imaging. The short optical distance between the bottom of the chip and the alveolus barrier, allows the use of standard air-dry objectives and confocal microscopy, enabling real-time visualization and quantitative analysis. Finally, to promote the interoperability and standardization among the existing lung on chip models, the 3DP-μLung was designed following a widely adopted architecture consisting of two microfluidic channels separated by a porous membrane (Huh et al, 2010, 2011, 2012)(Thacker et al, 2021; Bai et al, 2022; Van Riet et al, 2022; Mishra et al, 2023; Ektnitphong et al, 2024; Man et al, 2024). This configuration is also commonly used in available commercial solutions. In contrast to some lung on chip platforms, 3DP-μLung do not incorporate capacities for mimicking cyclic breathing motions (Huh et al, 2010; Huang et al, 2021; Sengupta et al, 2022; Luk et al, 2025). This design choice reflects our objective for the development of an open, accessible and cost-effective system that can be readily implemented in conventional academic laboratories.

Our study demonstrates that primary human alveolar epithelial cells can be used to reproducibly establish a functional alveolar barrier under physiological culture conditions. Because primary cells preserve key characteristics, including AT1 and AT-2 like cells and proliferative states of human adult lung tissue, their use offers several advantages over immortalized cell lines, particularly for controlling cell seeding density and proliferation dynamics. Because commercial primary cells can be expensive, we also optimized cell culture surface area of the device to reduce the number of primary cells and macrophages required, while still providing sufficient material for downstream analyses, such as qPCR quantitative image analysis. Recently, alternative approach based on iPSC or fetal stem cells demonstrated new opportunities for deriving autologous alveolar on chip (Kim et al, 2024; Luk et al, 2025) (Fonseca 2025). However, this approach typically requires significant expertise, costly or restricted cell access, and lengthy differentiation protocols that often exceed five to six weeks before infection can be initiated. (Wnorowski et al, 2019; Luk et al, 2025). In contrast, our approach can be established in less than 2 weeks, providing a rapid and practical platform for long term infection studies (up to 5 days post infection).

In this study, we investigated early host-pathogen interactions during *Mtb* infection. Our results indicate that Mtb infectivity within the alveolar barrier is primarily driven by macrophages. Strikingly, epithelial-only AoC models exhibit limited susceptibility to Mtb infection (H37Rv strain). In contrast, the integration of macrophage within the alveolar barrier markedly increases infection levels at the tissue scale, encompassing both macrophage and epithelial populations. This highlights the central role of macrophages as a permissive niche that facilitates the initiation, amplification, and dissemination of infection across the barrier during these early events. This also suggest that macrophages may act as vectors for epithelial colonization or alter the local microenvironment to favor bacterial uptake.

Moreover, we observed that the presence of macrophage promoted detectable intracellular bacterial replication over five days of infection. This supports the concept that macrophages provide a permissive niche for early *Mtb* growth, consistent with observations in traditional 2D cultures and other reports on lung on chip.(Cohen et al, 2018; Ahmad et al, 2022; Bo et al, 2023; Sankar & Mishra, 2023). Although epithelial infection remained relatively limited, our observations confirm that alveolar epithelial cells also participate in early infection events, consistent with previous reports (Ryndak & Laal, 2019; Moule & Cirillo, 2020; Sankar & Mishra, 2023; Olmo-Fontánez et al, 2024).

Interestingly, in these cell culture conditions, we observed events of macrophage transmigration toward the endothelial compartments. Notably, this occurred in the absence of detectable cell death or epithelial disruption, contrasting with several previous reports where infection led to barrier breakdown (Mishra et al, 2023). While these events were too sporadic for rigorous quantitative analysis (Fig. S3), this migratory behavior aligns with observations in other experimental systems (Thacker et al, 2020; Mishra et al, 2023; Ektnitphong et al, 2024). Thus, this observation may reflect a physiological phenotype of bacterial dissemination, contributing to the hematogenous spread of *Mtb* and the establishment of infection at distal sites. These findings highlight the unique capabilities of the organ-on-chip platform and the relevance of our model for studying early host-pathogen interactions at the tissue scale. This opens new opportunities to investigate the mechanisms of bacterial dissemination, a phenomenon difficult to observe in vivo and is poorly understood at the clinical level.

During this study, we also observed the formation of intracellular bacterial aggregates after five days of infection. However, our current imaging system did not allow us for the clear identification of cord-like structures under our experimental conditions. While this may be a technical limitation, the absence of such structures could also reflect significant differences in OOC maturation protocols, such as the implementation of air-liquid interface conditions and cyclic breathing motions. Unlike other studies, our protocol was optimized for rapid maturation of the alveolar barrier (7 days) and maintenance within a biosafety level 3 (BSL-3) environment. Furthermore, our decision to prioritize ease of use by excluding mechanical stretching forces precluded the investigation of mechanobiological mechanisms previously proposed to influence cording behavior. (Thacker et al, 2020; Mishra et al, 2023; Luk et al, 2025). Despite the absence of mechanical stretching forces, high-resolution imaging and quantitative analysis confirmed consistent *Mtb* infectivity, intracellular replication, and the formation of bacterial aggregates within both macrophages and epithelial cells.

In conclusion, we present a simple and cost-effective approach for modeling a human alveolus-on-chip platform that is fully compatible with BSL-3 laboratory environment for studying respiratory infection. By combining an open-source, accessible organ-on-chip (3DP-μLung chip) with high-resolution imaging, we have developed a robust tool for mimicking the physiologically relevant human alveolar environment. Our results demonstrate the platform’s capacity to provide deeper insights into the early events of Mtb infection at the human tissue scale.

## Materials and Methods

### Chip fabrication and assembly

The 3DP-µLung microdevice is composed by two distinct fluidically independent top and bottom channels separated horizontally by a polyethylene terephtalate (PET) porous membrane (culture surface area of 8.75 mm^2^), which simulate respectively the alveolar and vascular compartments, mimicking the alveolar biointerface and allow for cell-specific media.

3D printed molds for chips were designed using Fusion 360 (Autodesk) and printed from files uploaded in PreForm software (Formlabs) using a Form 3 printer (Formlabs) with Clear V4 resin (Formlabs) or Form 4 printer (Formlabs) with Clear V5 resin (Formlabs). Post treatment steps were performed previously described (Delannoy et al, 2025). Then chips were fabricated using the 3D printed molds and PDMS (Sylgard 184, Dow Corning) following soft lithography approach as previously described (Delannoy et al, 2025).

### Cell culture

Human primary alveolar epithelial cells (HPAEpiCs, Cat# H-6053, batch F040615Y72; Cell Biologics) were defrosted at P3 and expanded according to manufacturer’s instruction until P4 before use. Briefly, HPAEpiCs were grown in a treated culture flask pre-coated with gelatin-based coating solution (Cat# 6950; Cell Biologics) and incubated in complete growth medium (Cat# H-6621; Cell Biologics) for 5 days at 37°C, 5% CO^2^. Cells were detached with Accutase solution (Cat# A6964; Sigma-Aldrich) and frozen vials were prepared using at least 0.5 × 10^6^ cells within a solution of 90% heat-inactivated fetal bovine serum (FBS; Gibco) and 10% dimethyl sulfoxide (Sigma-Aldrich).

Human Lung endothelial cells (HULEC-5a, Cat# CRL-3244; ATCC) were grown in MCDB 131 Medium (Gibco) supplemented with 10% FBS, 10 mM L-Glutamine (Gibco), 1 µg/mL Hydrocortisone (Cat# H0888; Sigma-Aldrich), 10 ng/mL recombinant human Epidermal Growth Factor (Cat# AF-100-15; PeproTech) and 1% Penicillin/Streptomycin (Cat# 15140-122; Gibco) in 5% CO_2_ at 37°C.

Cells vials were verified mycoplasma-free (MycoAlert Mycoplasma Detection Kit, Lonza).

### Isolation, differentiation and labelling of human monocyte-derived macrophages for AoC co-cultured with human macrophages (M-AoC)

Peripheral blood mononuclear cells (PBMCs) were obtained from buffy coats from three anonymized donors (provided by Regional French Blood Company, France). PBMCs were isolated using a Ficoll separation procedure, as previously described (Deboosere et al, 2021). CD14^+^ monocytes were isolated by magnetic cell separation, using cell labeling with magnetic human CD14 microbeads (Cat# 130-050-201; Miltenyi Biotec) and column-based separation (Cat# 130-042-401; Miltenyi Biotec), according to the manufacturer’s instructions. Isolated monocytes were cryopreserved, as previously above.

One week prior to seeding the human macrophages in the AoC devices, a cryopreserved aliquot of monocytes was thawed and cultured in RPMI medium (Gibco) supplemented with 10% FBS and 1% Penicillin/Streptomycin and differentiated for 7 days with 40 ng/mL recombinant human macrophage-colony stimulating factor protein (hM-CSF, Cat# 130-096-491; Miltenyi Biotec). To vizualize macrophages co-cultured with HPAEpiCs, CellTrace CFSE solution at a final concentration of 10 µM (Cat# C34570; Invitrogen) was used as described in the manufacturer’s instructions. After centrifugation, cells were resuspended at 2.5 × 10^5^ cells/mL in complete epithelial growth medium.

### Human Alveolus-on-Chip seeding and culture

Both channels of 3DP-µLung were washed with Dulbecco’s Phosphate-Buffered Saline no calcium, no magnesium (DPBS -/-; Gibco) and coated overnight at 37°C with 200 μg/mL Collagen IV (Cat# C5533; Sigma-Aldrich) and 30 μg/mL Fibronectin from human plasma (Cat# 356008; Corning) diluted in cold DPBS -/-. The next day (day 0), both channels were rinsed with warm complete endothelial cell culture and HULEC-5a cells seeded in the bottom channel at 1 × 10^7^ cells/mL. Chips were flipped and placed on chip custom made cradles for 4 h at 37°C, 5% CO_2_. Then, channels were rinsed twice with 100 µL of warm complete endothelial cell culture. Chips were maintained overnight under static conditions. On day 1, HPAEpiC were thawed and seeded at 2.5 × 10^6^ cells/ mL in the top channel. Chips were placed right side up for 4 h at 37°C, washed and incubated statically overnight. On day 2, confluent AoC are connected to microfluidic circuit actuated by a peristaltic pump (Ismatec, Switzerland), regulating media flow at 60 µL/h in both channels. Apical and basal channels were perfused with cell-specific growth media, and epithelial medium was supplemented with 1 μM dexamethasone (Cat# D4902; Sigma-Aldrich) to promote surfactant production and formation of tight junctions, as reported in previous lung-on-chip studies (Huh et al, 2010; Thacker et al, 2021; Mishra et al, 2023; Ektnitphong et al, 2024). Chips were cultured under flow conditions for 4 days at 37°C, 5% CO_2_. On day 6, the apical channel medium is removed, to establish an air-liquid interface (ALI). The upper channel was maintained in air static condition by introducing emptied filtered tips in the inlet and outlet of the top channel.

On day day 7, approximately 5 × 10^3^ of differentiated human CD14+ peripheral blood-derived macrophage cells, labeled with CFSE (corresponding to a ratio of 10% as compared to epithelial cells) and seeded (chip defined as M-AoC). Macrophages were allowed to adhere in static condition overnight at 37°C, 5% CO_2_. On day 8, chips were washed with warm epithelial cell-specific medium and kept in LLI.

### Mycobacterium tuberculosis infection

Recombinant strains of Ds-Red-expressing *Mtb,* H37Rv-RFP (H37Rv-DsRed) and H37Ra-RFP (H37Ra-DsRed), were cultured in Middlebrook 7H9 medium (Difco) supplemented with 10% oleic acid-albumin-dextrosecatalase (Difco), 0.2% glycerol (Euromedex), 0.05% Tween 80 (Sigma-Aldrich), and 25 mg/mL kanamycin (Sigma-Aldrich) for up to 14 days before being used for the infection assay, to reach the exponential phase of bacterial growth, as previously described (Deboosere et al, 2021).

*Mtb* culture grown to exponential phase (OD600 of 0.5) was centrifuged at 5000 g for 5 minutes at RT. After supernatant removal, the cell pellet was washed two times with DPBS -/- and resuspended in complete epithelial cell media. Clumped mycobacteria were removed by centrifugation at 800 rpm for 2 min. And a single cell suspension was obtained via filtration through a 10 µm cell pluristrainer (Cat# 43-50010-01; pluriSelect). The single-bacteria suspension was diluted in epithelial media to obtain a bacteria suspension at 5. 10^6^ bacteria/mL.

We infected eight days old AoC or M-AoC at an estimated multiplicity of infection (MOI) of 5. Chips were incubated in static condition for 4 h at 37°C, 5% CO_2_ to allow *Mtb* infection of the alveolar epithelium. Then, both channels were washed twice with 100 µL of cell medium and treated with amikacin (Sigma-Aldrich) for 1 h at 37°C to kill remaining extracellular bacteria. Followed by two additional washes, AoC were maintained in liquid-liquid interface (LLI) and in static condition until 5 dpi. For safety precautions from our BSL3 lab, fluidic regulation could not be operated.

### Immunofluorescence

Prior fixation in 4% paraformaldehyde solution (PFA; Biovalley) in DPBS with Ca^2+^ and Mg^2+^ (DPBS +/+; Gibco), DPBS +/+ were flushed in both channels. Cells were fixed for 30 min at room temperature (RT) in non-infectious condition and overnight in infectious condition, followed by three washes with DPBS +/+ for both conditions. Then autofluorescence was quenched with 50 mM NH_4_Cl in DPBS +/+ for 30 min at RT and rinsed twice with DPBS +/+. AoC were stored at 4°C in DPBS+/+ until further processing.

For immunostaining, AoCs were first permeabilized in DPBS +/+, 0.5% BSA (Sigma-Aldrich), 0.2% Triton X-100 (Sigma-Aldrich) (permeabilization buffer) for 15 min at room temperature (RT) and rinsed twice with DPBS +/+, 0.5% BSA, 0.2% FBS (washing solution). Cells were incubated with a blocking solution DPBS +/+, 5% BSA, 2% FBS for 1h at RT, followed by two washes. For cell characterization, AoC was incubated with the primary antibodies HT1 (Cat# TB-29 AHT1-56; Terrace Biotech) and HT2 (Cat# TB-27 AHT2-280; Terrace Biotech) or VE-Cadherin (Cat# 348502; BioLegend), diluted in permeabilization buffer overnight at 4°C. Then chips were washed 3 times for 5 min with washing solution and incubated for 2h at RT with the specific secondary antibodies diluted in permeabilization buffer. F-actin was stained using Phalloidin-coupled to Alexa 647 (Cat# A22287; Thermofisher) concurrently with 10 µg/mL DAPI (Cat# D1306; Thermofisher) in permeabilization buffer for 1h at RT, rinsed twice for 5 min with DPBS +/+ and stored at 4 °C until observation. For M-AoC experiments, an additional immunostaining was performed using CD68 antibody (Cat# sc-20060; Santa-Cruz). Overnight incubation was performed with CD68 antibody diluted in permeabilization solution at 4°C. The following day, samples were flushed three times extensively with washing solution and rinsed at least 2 times in DPBS -/- and immediately imaged. For transversal section imaging, chips were cut in 1 mm thin segments using a razor blade or vibratome. Transversal sections were kept at 4°C in a 24-well plate containing DPBS +/+ and mounted with DPBS +/+ on glass coverslip prior to imaging. Primary and secondary antibodies used in this study are listed in the following table (Table 1).

**Table 1.** List of antibodies used for immunofluorescence staining.

| ANTIBODY | HOST/ISOTYPE | MANUFACT. | CAT # | DIL. |
| --- | --- | --- | --- | --- |
| **Primary antibody** |  |  |  |  |
| Anti-human CD144 (VE-Cadherin) | Mouse / IgG2a, k | BioLegend | 348501 | 1/50 |
| Anti HT1-56 (AT-I) | Mouse / IgG1 | Terrace Biotech | TB-29 AHT1-56 | 1:10 |
| Anti HT2-280 (AT-II) | Mouse / IgM | Terrace Biotech | TB-27 AHT2-280 | 1:10 |
| **Secondary antibody** |  |  |  |  |
| Anti Mouse IgG (H+L) Alexa Fluor 488 | Goat | Invitrogen | A32723 | 1:500 |
| Anti Mouse IgG1 recombinant VHH Alexa Fluor 568 | Alpaca | Chromotek | sms1AF568-1 | 1:500 |
| Affinipure Anti Mouse IgM μchain specific Alexa Fluor 488 | Donkey | Jackson ImmunoResearch | 715-545-140 | 1:500 |
| **Other** |  |  |  |  |
| Phalloidin Alexa Fluor 647 |  | Invitrogen | A22287 | 1:500 |
| CD68 Alexa Fluor 488 |  | Santa Cruz | sc-20060 AF488 | 1:50 |

### Live-dead assay

Cell viability was assessed using the LIVE/DEAD™ Viability/Cytotoxicity Kit (ThermoFisher Scientific) according to the manufacturer’s protocol. Briefly, a staining solution was freshly prepared by diluting calcein-AM (2 μM final concentration) and ethidium homodimer-1 (4 μM final concentration) in PBS. Live non-infected AoC were incubated with the staining solution for 40 minutes at 37 °C. Following incubation, chip channels were washed extensively with culture medium and imaged directly using a spinning disk microscope (Nikon - Gataca Systems). Quantification was performed by analyzing images using ImageJ software. The percentage of viable cells was calculated as the ratio of calcein-positive cells to the total number of cells (calcein-positive + ethidium homodimer-positive).

### qRT-PCR

Channels were washed with DPBS +/+ into the channel inlets, followed by the injection of 100 μL RNA lysis buffer (RNeasy Plus Micro Kit, Cat# 74034; Qiagen) into top channel for 5 min incubation time. Then cell lysate was quickly homogenized using a micropipette plunger. The procedure was repeated 3 times to collect a total volume of 300 µL of lysates. Samples were stored at −80 °C until RNA analysis.

After determining RNA concentrations by spectrophotometry, 50–500 ng of total RNA was used for cDNA synthesis. Reverse transcription was conducted using the High-Capacity cDNA Reverse Transcription Kit (Cat# 4368814; Applied Biosystems). Quantitative real-time PCR was performed using the Takyon™ Low Rox SYBR MasterMix dTTP Blue (Cat# UF-LSMT-B0101; Eurogentec) with 300 nM primers and 1:12.5 prediluted cDNA. The specificity of primers was confirmed by melting curve analysis and gel electrophoresis. qPCR was performed on a QuantStudio 3 System (Applied Biosystems). Relative RNA level was quantified using the ΔΔCt method and normalized to the endogenous control GAPDH. Specific primers (Sigma-Aldrich) used for the qRT–PCR were previously used (Thacker et al, 2021) and listed in Table 2.

**Table 2.**
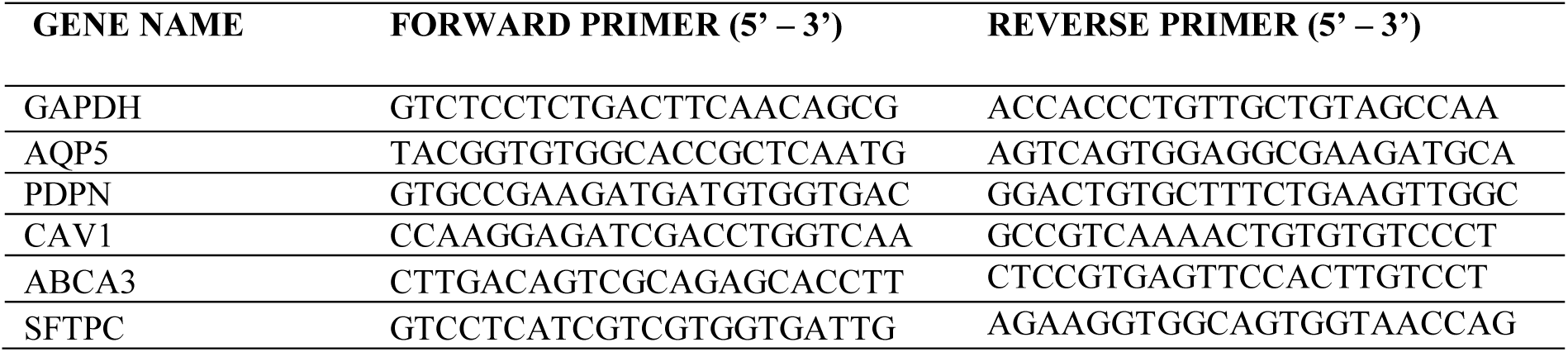
List of forward and reverse primers used for qRT-PCR.

### Permeability assay

AoC maturation was evaluated by microscopy and by the permeability assay on day 7 and day 13. Chips that did not passed this quality control due to non-confluent cell areas were used as controls and named « unfunctional AoC ». All chips were evaluated using 10 µg/mL FITC-dextran 70 kDa (Cat# D1823; Invitrogen) diluted in cell culture media and flowed overnight at 60 µL/h within the top channel within the incubator at 37 °C, 5% CO_2_. Samples from the top and bottom channel outlets were collected and fluorescence levels were measured a VictorX3 multilabel plate reader (PerkinElmer) with excitation wavelength of 485 nm and an emission wavelength of 535 nm. The apparent permeability coefficient (Papp) was calculated based on the final fluorescence values using the following simplified formula:

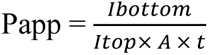

where Ibottom and Itop are the fluorescence intensities measured at the end of the experiment in the top and bottom channels, respectively, A is the surface area of the permeable membrane (cm^2^), and t is the incubation time (s). This approach assumes a linear transfer of dextran during the incubation period.

### Microscopy imaging

The maturation of uninfected alive AoC was followed by phase contrast imaging with an inverted microscope (Eclipse Ts2; Nikon) equipped with a camera (Digital Sight 1000; Nikon). The lower and upper channels were controlled with 4X and 20X objectives (Nikon Plan Apochromat Lambda D 4x 0.20 N.A and 20x 0.4 N.A). For quantitative analysis, all images were acquired using a spinning disk microscope (Nikon - Gataca Systems) equipped with a CSU-W1 confocal scanner unit and Live-SR module. A Z-series of optical sections was acquired covering approximately 100 μm and casting the endothelial cell layer, the porous membrane and the epithelial cell layer. First, for controlling the alveolar barrier integrity, a full scan of the channel was performed using an objective 10× dry CFI Plan Fluor (0.30 NA; Nikon) with 405 and 647 nm lasers or 405, 488 and 561 nm lasers. Then, for characterizing and quantifying infection experiments of the alveolar barrier, fluorescence images from AoC were taken using the objective 20× dry CFI Plan Apo LBDA (0.75 NA; Nikon). Three regions of channel area were acquired along the channel.

### Image analysis

Image J software (v1.54m, National Institutes of Health) and Imaris 3D microscopy image analysis software (v10.1.1, Oxford Instruments) were used for imaging analysis. Images were adjusted for brightness, color balance, and/or contrast uniformly across all pixels, as necessary.

For infection experiments, 3 fields of view along the chip were used and fluorescence images were firstly processed and improved using a median filter. Then, nuclei (Hoechst-labeling) and H37Rv-DsRed bacteria were detected using a spot detection module based on fluorescence intensities. Bacterial area was measured in pixels. The bacterial replication represents the total bacterial area per region of interest (ROI). Infection rate was determined as the total number of detected bacteria spots for each condition by the total of cells detected. The bacterial growth was quantified as the total intracellular bacterial area per ROI.

### Statistical analysis

Infection experiments were repeated six times, with three biological replicates. Data are displayed as mean values ± standard deviation (SD) unless otherwise noted in the figure legends. Graphing and statistical comparison of the data were performed using Prism 10.4.1 (GraphPad Prism Software). Two-group comparison was assessed using the unpaired t test for the permeability assay; and non-parametric Mann-Whitney test was used to compared unpaired data in infection experiments. P values < 0.05 were statistically significant.

## Data Availability

The authors confirm that the data supporting the findings of this study are available within the article

## Acknowledgements

We thank Cecile Lecoeur and the Biostatistics Core Facility of the CIIL for the guidance on statistics. We thank BICeL - PLBS -UMS 2024 -US 14 and Olivier Molendi-Coste for the guidance on flow cytometry analysis. We thank members of MoHMI team for comments and discussion on the manuscript; Cyril Gaudin, Eik Hoffmann, Arnaud Machelart, Amine Pochet, Joan Fine for assistance with BSL-3 laboratory experiments.

This work was supported by Atip-Avenir funding, MEL “Accueil de talents” funding, INSERM, the Agence Nationale de la Recherche (ANR-18-JAM2-0002, 20-AMRA-0005, and ANR-22-CE18-0021-01), the Agence Nationale de Recherche sur le SIDA et les hépatites virales - Maladies Infectieuses Emergentes (ANRS-MIE No. ANRS-0521 and ANRS-0655), and the Institut Pasteur (PTR 430-21 and PTR 22-16). This project has received funding from the Innovative Medicines Initiative 2 Joint Undertaking (JU) under grant agreement No. 853989 (ERA4TB). French government under the France-2030 program, the University of Lille and the Lille European Metropolis (MEL) are thanked for their funding and support for the project R-CDP-24-007-MOSAIC granted to AG. Y.D. was the recipient of a fellowship from the Agence Régionale de Santé (ARS).

The funders had no role in study design, data collection and analysis, decision to publish, or preparation of the manuscript. The authors declare that they have no competing interests. A figure was created using the software BioRender, licence to Yoël Dagan.

## Author contributions

N.D. and Y.D. designed, performed, and analyzed most of the experiments; E.D. designed the microfluidic chip; A.B. fabricated the microfluidic chips; S.S. and E.W. participated in image analysis. R.S. contributed to funding acquisition. A.G. and P.B. supervised the work, secured funding, participated in the design and analysis of most experiments, and wrote the manuscript with N.D.

## Supplemental Figures

**Figure S1.**
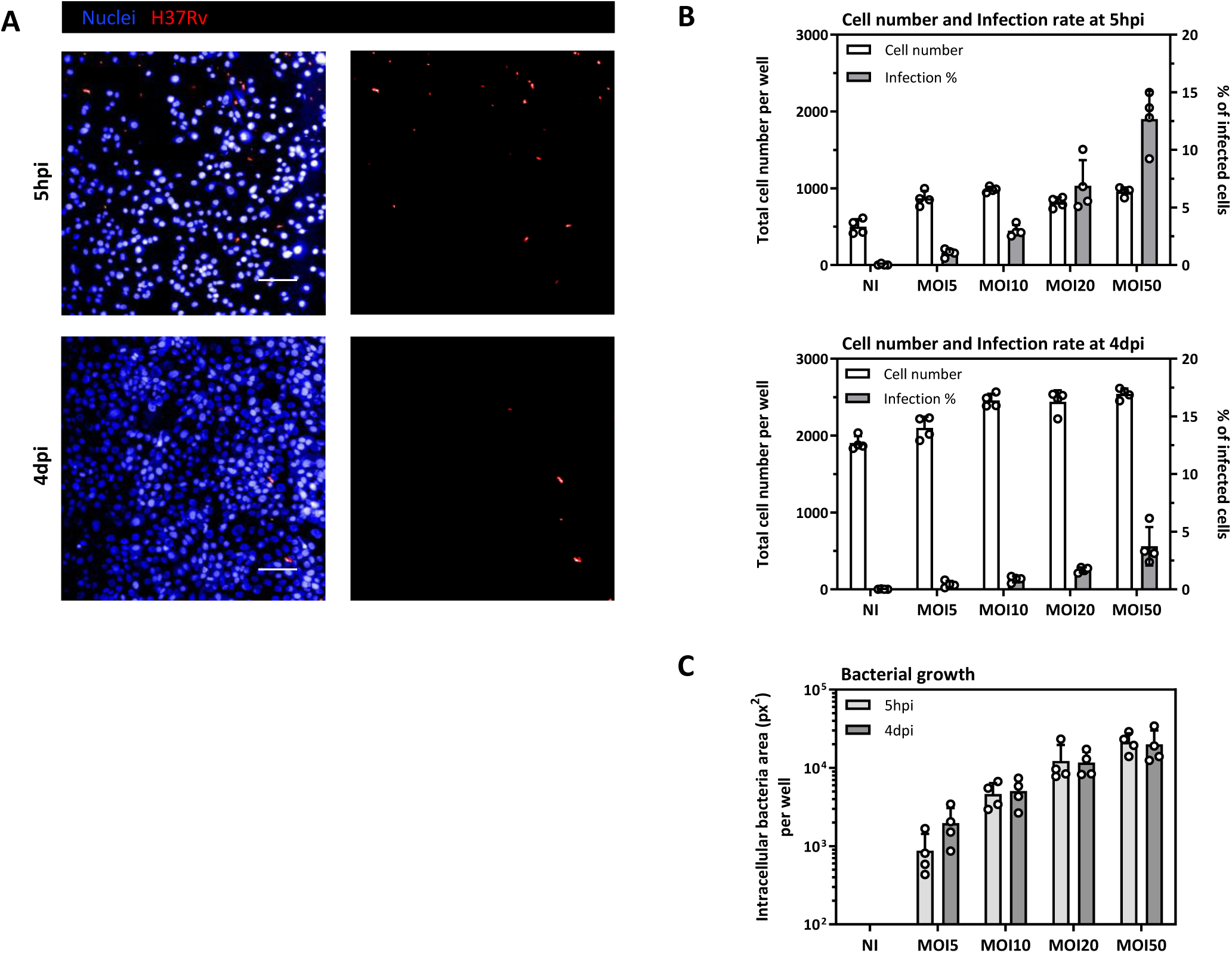
*Mtb* infection in 2D-culture model of HPAEpiC in 384-well plate. **(A)** Typical images of infected HPAEpiC in 2D-culture model on day 0 (5hpi) and day 4 (4dpi), used for the determination of *Mtb* uptake and intracellular growth inside epithelial cells. HPAEpiC were grown in 384-well plates, infected with *Mtb* H37Rv-DsRed and imaged with an automated confocal microscopy (InCell6005-GE Healthcare). Shown are representative images of infected HPAEpiC with MOI 10 at 5hpi (uptake on D0) and 4dpi (bacterial growth); Nuclei (Blue) and H37Rv (Red). Scale bar= 100 μm. **(B, C)** Quantitative image analysis of infected HPAEpiC in 2D-culture model at 5hpi and 4dpi. **(B)** Total cell numbers and percentage of cell infection and **(C)** bacterial growth. Shown are mean of four analyzed wells per condition of two representative experiment.

**Figure S2.**
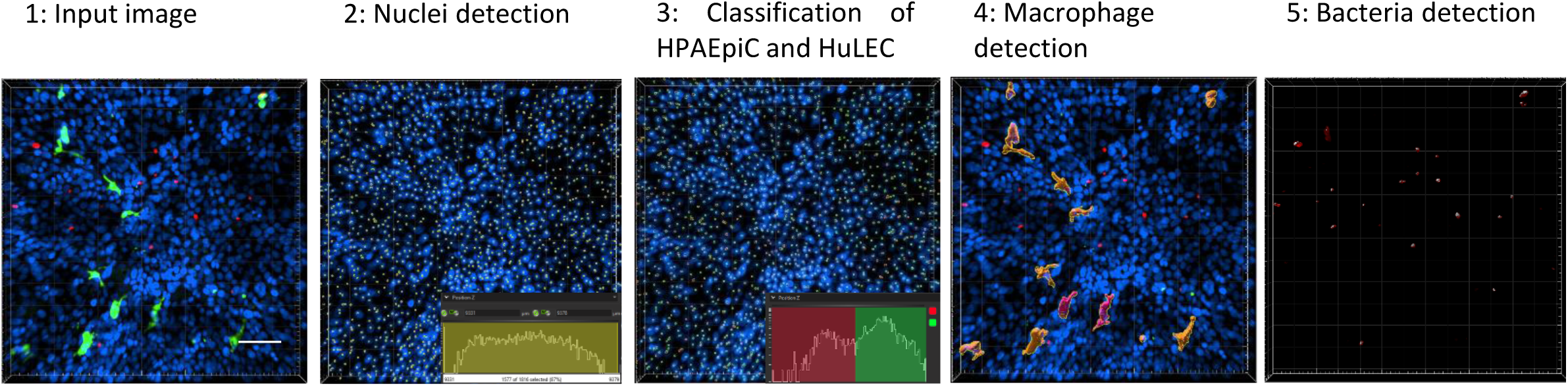
Typical images of controls and the corresponding detection used for the determination of *Mtb* infection in Alveolus-on-Chip. Images of infected AoC were acquired by confocal microscopy (Spinning-Disk-Nikon - Gataca Systems) and a quantification of the infection was performed using 3D image analysis software (IMARIS v10, Oxford Instruments). Detection tools were applied to input images to detect nuclei labeled by Hoechst 33342 (blue), F-actin with Alexa Fluor 657 Phalloidin dye (gray), macrophages with anti CD68 antibody conjugated to Alexa Fluor 488 (green), and the DsRed signal of H37Rv-DsRed (red), to determine cell numbers and macrophage and bacteria total volumes.

**Figure S3.**
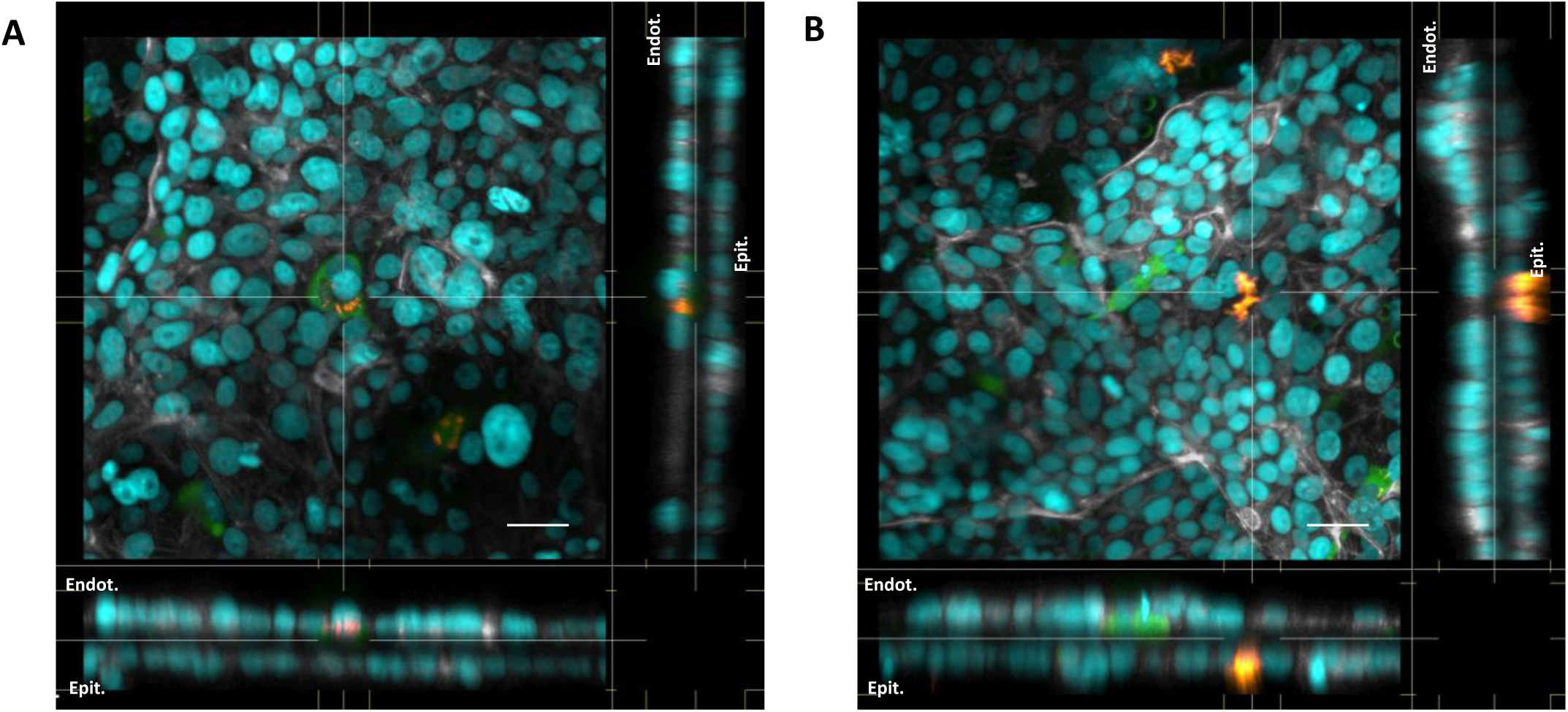
Intracellular *Mtb* H37Rv in Alveolus-on-Chip co-cultured with human macrophages (M-AoC). **(A, B)** Immunofluorescence imaging of infected M-AoC at 5dpi, using the Nikon Spatial Array Confocal (NSPARC) detector on the AX system (Nikon). **(A, B)** Maximum intensity xy projections and extended orthogonal xz and yz cross-sections from two fields of infected M-AoC. Nuclei (Cyan), H37Rv (Yellow), Actin (Gray) and macrophages (Green). Scale bar= 30 μm.

**Video S1.**
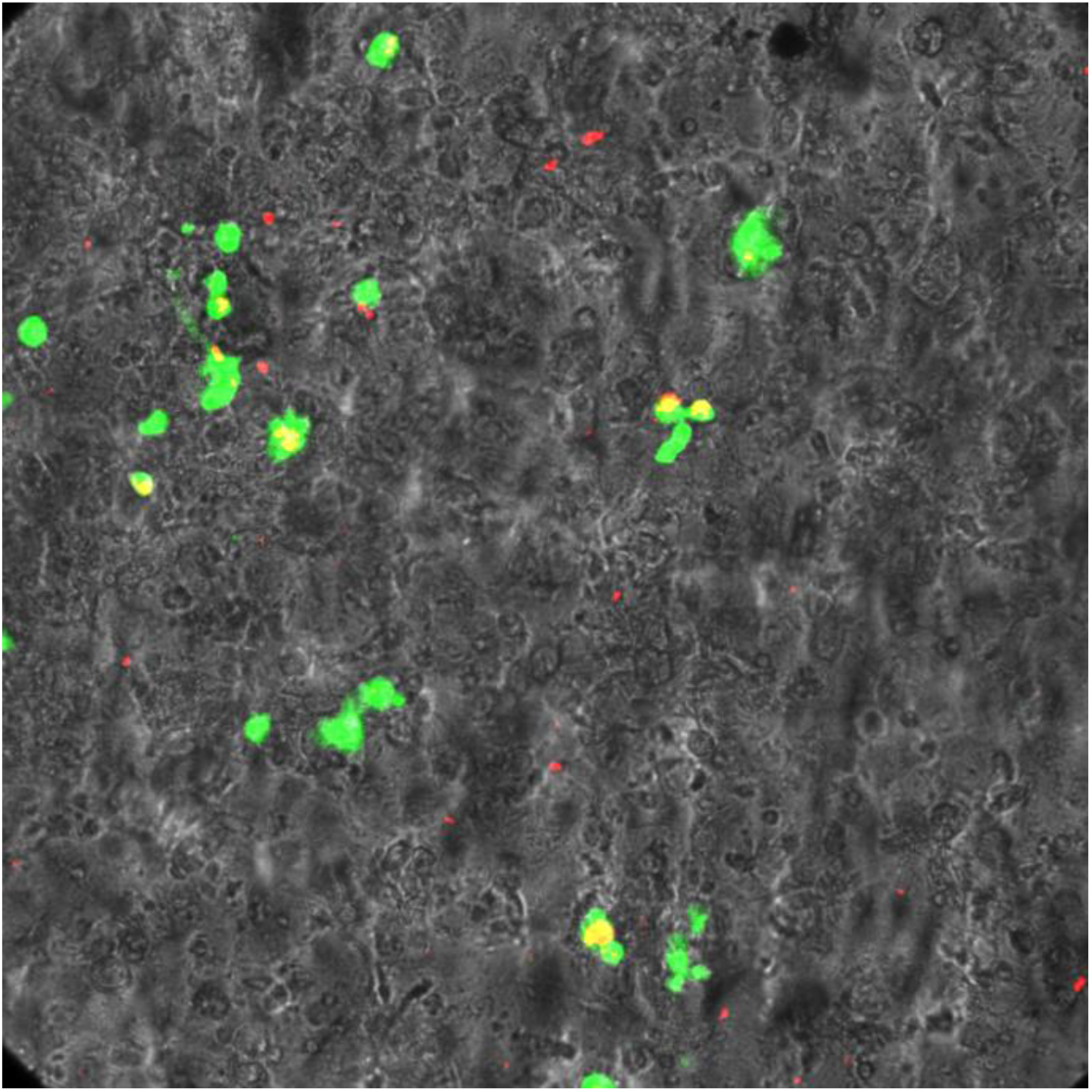
Time-lapse microscopy of Alveolus-on-Chip co-cultured with human macrophages (M-AoC) infected with *Mtb* H37Ra-DsRed. Live-cell imaging was run over 17 hours post-infection on a confocal microscopy (Spinning-Disk-Nikon - Gataca Systems). An AoC co-cultured with CFSE-labeled human macrophages was infected with *Mtb* H37Ra-DsRed at MOI 5. Images were taken every 30 minutes, highlighting th e intracellular localization of H37Ra within macrophages. H37Ra (Red) and macrophages (Green). Field of view is 4.3×10^5^ µm^3^. Scale bar=100μm.

